# Endogenous zebrafish proneural Cre drivers generated by CRISPR/Cas9 short homology directed targeted integration

**DOI:** 10.1101/2020.07.21.214452

**Authors:** Maira P. Almeida, Jordan M. Welker, Stephen C. Ekker, Karl J. Clark, Jeffrey J. Essner, Maura McGrail

**Author notes:** Corresponding author: Maura McGrail 2213 Pammel Dr., 3005 ATRB, Iowa State University, Ames, Iowa, 50011. NIH R24OD020166 (MM, JJE, SCE, KJC), CNPq Brazilian National Council for Scientific and Technological Development (MPA).

## Abstract

The Cre/*lox* recombinase system has been widely used for spatiotemporal control of gene expression in animal model systems, however, efficient methods to isolate zebrafish Cre drivers that reliably recapitulate endogenous gene expression patterns are needed. Here, we apply CRISPR/Cas9 targeting to integrate a 2A-Cre recombinase transgene with 48bp homology arms into proneural genes *ascl1b*, *olig2* and *neurod1*. We observed high rates of germline transmission ranging from 10%-100% (2/20 *olig2*; 1/5 *neurod1*; 3/3 *ascl1b*). The transgenic lines *Tg*(*ascl1b-2A-Cre)^is75^*, *Tg*(*olig2-2A-Cre)^is76^*, and *Tg*(*neurod1-2A-Cre)^is77^* expressed functional Cre recombinase in the expected proneural cell populations. The results demonstrate Cre recombinase expression is driven by the native promoter and regulatory elements of the targeted genes. This approach provides a straightforward, efficient, and cost-effective method to generate cell type specific zebrafish Cre drivers whose spatial and temporal restricted expression mimics endogenous genes, surmounting the challenges associated with promoter BAC cloning and transposon mediated transgenesis.

## Introduction

The Cre/*lox* recombinase system has been widely used in zebrafish for spatiotemporal control of gene expression and lineage tracing (Hans et al., 2009; Kirchgeorg et al., 2018; Mosimann et al., 2011; Sanchez-Iranzo et al., 2018). The effectiveness of the system is dependent on drivers that provide precise spatial and temporal Cre expression, and recent technical advances combining multiple recombinases have increased the ability to restrict activity to defined cell populations (Liu et al., 2020; Poulin et al., 2020). However, the approach is still limited by methods used to isolate zebrafish transgenics which are susceptible to position effect and multigenerational silencing. An improved method to isolate recombinase transgenics that recapitulate endogenous gene expression patterns would increase the robustness and reproducibility of Cre/lox lineage tracing and gene function studies.

In organisms such as zebrafish that lack embryonic stem cell methodology, Tol2-mediated transgenesis (Balciunas et al., 2006; Kawakami et al., 2000) has traditionally been used to isolate Cre drivers (Carney and Mosimann, 2018). In one approach, a gene promoter is cloned or BAC-engineered into a promoter-Cre fusion inside a Tol2 vector (Forster et al., 2017; Mosimann et al., 2011). Alternative strategies using Tol2 transgenesis exploit the nearly unbiased, genome wide random integration of the transposon to isolate novel cell type and tissue-specific drivers. Enhancer trap Tol2 vectors contain a minimal promoter-Cre-2A-reporter cassette and are designed to drive expression under the control of local enhancer elements at the integration site (Marquart et al., 2015; Tabor et al., 2019). Tol2 vectors engineered with a splice acceptor-mCherry-2A-Cre-ERT2 gene trap are driven by endogenous promoter and regulatory elements after in-frame integration within a gene (Jungke et al., 2015). Although each Tol2-based method allows for efficient recovery of transgenics, difficulties associated with defining and cloning a complete promoter and associated regulatory elements, combined with multicopy transgene integration, position effects and gene silencing, limits the effectiveness of creating drivers with consistent restricted expression patterns (Hans et al., 2009; Mukherjee and Liao, 2018; Yang and Gong, 2005).

Recent advances in gene editing with engineered site-specific nucleases may solve a number of the limitations associated with Tol2 transgenic approaches used to isolate zebrafish transgenic Cre drivers. The ability of TALENs and CRISPR/Cas9 to direct a double stand break to a specific location in the genome and generate single stranded overhangs for DNA repair has been exploited to introduce exogenous DNA at the target site via non-homologous end-joining (NHEJ) and homology directed repair (HDR) pathways (Bedell et al., 2012; Beumer et al., 2008; Carlson et al., 2012; Suzuki et al., 2016; Yang et al., 2013). Early studies in zebrafish demonstrated the addition of complementary homology arms flanking a donor transgene significantly increased the frequency and precision of targeted integration (Hisano et al., 2015; Hoshijima et al., 2016; Shin et al., 2014). One report in zebrafish described using CRISPR and 1 kb of homology cloned in front of a targeting cassette to integrate Cre-ERT2 upstream of the *otx2* translation initiation codon (Kesavan et al., 2018). However, the efficiency of precision integration using this approach is not known.

To streamline isolation of zebrafish Cre drivers for specific cell lineages and cell types, we applied our recently reported method for CRISPR/Cas9 short homology directed targeted integration (Wierson et al., 2020) to generate Cre transgenic lines. As proof of principle we targeted a 2A-Cre cassette in frame into a coding exon in the proneural transcription factor genes *ascl1, olig2* and *neurod1* that define neural stem, progenitor and post-mitotic cell lineages (Aprea et al., 2014; Bertrand et al., 2002; Hevner et al., 2006; Kim et al., 2011; Takebayashi et al., 2002; Wilkinson et al., 2013; Zhou and Anderson, 2002). We chose to target *ascl1b* and *olig2* since both are differentially expressed in highly aggressive pediatric brain cancers characterized by embryonal-like, poorly differentiated tumors (Picard et al., 2012; Sturm et al., 2016), consistent with their role in progenitor cell function during neural development. We and others had shown previously *ascl1b* and *olig2* are overexpressed in zebrafish embryonal brain tumor models, reinforcing the conservation of molecular mechanisms driving neuroectodermal tumor types (Modzelewska et al., 2016; Schultz et al., 2018). Together with *neurod1*, a marker of early neural commitment and differentiation, *ascl1b* and *olig2* are good candidates to generate Cre drivers that would be useful for functional studies in neurogenesis and brain tumor pathogenesis. The 2A-Cre targeting vector used for integration is part of our pPRISM vector series that contains cassettes with linked fluorescent secondary marker for allele tracking (Welker et al., in preparation). Recovery of precise 2A-Cre integration alleles was highly efficient, with frequencies ranging from 10-100%, similar to other cargos as reported previously (Wierson et al., 2020). As expected, the expression of functional Cre recombinase was restricted to cell populations defined by *ascl1b, olig2* and *neurod1*. Together, our results demonstrate CRISPR/Cas9 directed targeted integration is an efficient and robust method for isolating endogenous Cre drivers that reflect the endogenous promoter activity of the targeted genes.

## Results and Discussion

### CRISPR/Cas9 mediated Knock-in of 2A-Cre into *ascl1b, olig2*, and *neurod1*

To generate proneural Cre drivers we integrated a 2A-Cre recombinase cDNA cassette in frame into a coding exon of the zebrafish *ascl1b, olig2* and *neurod1* genes (Figure 1). Our strategy for CRISPR/Cas9 precision targeted integration uses short homology arms flanking a targeting cassette to likely drive integration by homology mediated end joining (HMEJ) (Wierson et al., 2020). The 2A-Cre targeting construct contains a secondary fluorescent marker cassette with the *γ-crystallin* (*γ-cry*) promoter driving EGFP expression (Figure 1A). Efficient targeted integration after injection into embryos is driven by *in vivo* liberation of the homology arms flanking the cassette, which are released by Cas9 induced double strand breaks at the universal gRNA (UgRNA) sites in the vector (Figure 1A).

**Figure 1.**
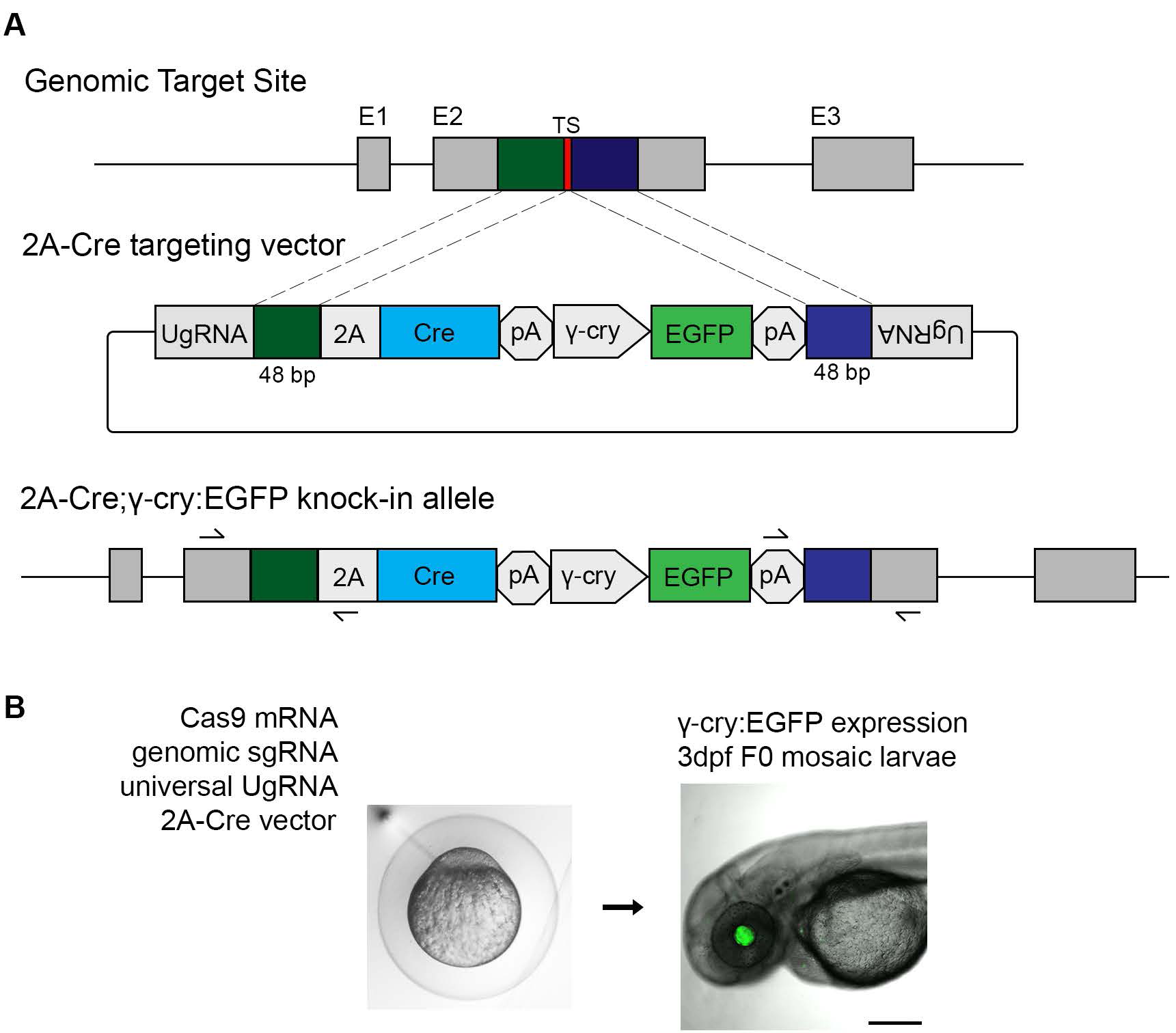
CRISPR/Cas9 short homology directed targeted integration strategy for efficient recovery of Cre knock-in alleles. (A) Schematic of the genomic target site, Cre donor vector, and final knock in allele. A CRISPR sgRNA (red) was chosen in the coding sequence. 48 bp upstream (dark green) and 48 bp downstream (dark blue) of the cut site are included in the donor vector as homology arms. The donor vector has UgRNA sequences flanking the homology arms for *in vivo* liberation of the knock in cassette. The knock-in cassette contains an in frame 2A-Cre for expression of Cre recombinase, and a secondary marker driving *γ-cry:EGFP* expression in the lens. Arrows indicate PCR primers for 5’ and 3’ genome/vector junction analysis. (B) 1 cell stage zebrafish embryos are injected with Cas9 mRNA (300 pg), genomic sgRNA (25 pg), UgRNA (25 pg), and 2A-Cre targeting vector (10 pg). At 3 days post fertilization, embryos expressing GFP in the lens are selected and raised to adulthood. *TS*, genomic sgRNA target site; *UgRNA*, universal short guide RNA target site; *γ-cry*, gamma crystallin promoter; *GFP*, green fluorescent protein; *pA*, transcription termination and polyadenylation sequence.

To isolate the 2A-Cre transgenic lines, sgRNAs were designed to CRISPR/Cas9 sites in exon 1 of *ascl1b* and exon 2 of *olig2* and *neurod1* (Table 1). The sgRNA mutagenesis efficiency was tested by co-injection with Cas9 mRNA into 1-cell zebrafish embryos, followed by PCR amplification of the targeted exon and analysis of heteroduplex formation in the PCR product by gel electrophoresis. After confirmation of efficient mutagenesis, 48 bp of sequence on either side of the genomic target Cas9 was used to design and clone 5’ and 3’ homology arms into the 2A-Cre targeting vector. Zebrafish embryos were injected at the 1-cell stage with the genomic target site sgRNA, the UgRNA, Cas9 mRNA, and the 48 bp homology arm 2A-Cre targeting vector. At 3 days post-fertilization (dpf), larvae were screened for expression of the lens specific γcry:EGFP secondary marker (Figure 1B), and positive larvae selected to raise to adulthood to test for germline transmission. Targeting of *ascl1b, olig2* and *neurod1* resulted in 58%, 48% and 37%, respectively, of F0 injected embryos showing EGFP expression in the lens (Table 1), demonstrating the presence of the injected targeting vector. Evidence of somatic targeted integration at the genomic site was confirmed by junction fragment PCR on injected embryo.

**Table 1.** Efficient recovery of endogenous neurogenic Cre driver lines by 2A-Cre-γcry:EGFP targeted integration

| **Neurogenic Gene** | ***ascl1b*** | ***olig2*** | ***neurod1*** |
| --- | --- | --- | --- |
| Targeted Exon | 1 | 2 | 2 |
| 5' and 3' Homology Arms (bp/bp) | 48/48 | 48/48 | 48/48 |
| F0 embryos expressing secondary marker | 58% (61/105) | 48% (52/109) | 37% (21/57) |
| F0 adults transmitting secondary marker | 100% (3/3) | 10% (2/20) | 20% (1/5) |
| F0 adults transmitting precise integration allele | 33% (1/3) | 5% (1/20) | 20% (1/5) |

To test whether somatic integration of the 2A-Cre cassette led to expression of functional Cre recombinase in the expected neural cell populations, we injected the genomic sgRNA, UgRNA, Cas9 mRNA, and the donor vector into embryos from the transgenic ubi:Switch floxed reporter line *Tg(ubi:loxP-EGFP-loxP-mCherry)* (Mosimann *et al*., 2011).

Injected F0 mosaic *Tg(ubi:loxP-EGFP-loxP-mCherry*) embryos showed a switch from EGFP to mCherry expression in cells throughout the brain and retina, while EGFP expression remained in cells outside of the central nervous system (Supplementary Figure 1). These results suggested that Cre expression was controlled by the endogenous regulatory elements of the targeted proneural genes, leading to recombination in the specific neural progenitor populations defined by those genes. Furthermore, the results suggest that the expressed Cre has recombinase activity that leads to recombination at *loxP* sites and excision of the floxed *EGFP* cassette. All neural progeny descended from the neural progenitor populations inherit the recombination event and express mCherry. The observed recombination in F0 injected animals indicated on-target integration of the 2A-Cre cassette was relatively efficient. Together, these results suggested targeted integration would be an effective method to generate endogenous Cre drivers that can promote spatially restricted cell-type specific Cre-mediated recombination.

To isolate stable germline 2A-Cre knock-in alleles, adult F0s were outcrossed to wildtype WIK to identify individuals transmitting the γcry:EGFP secondary marker to their progeny. We first measured the frequency of germline transmission of the γcry:EGFP secondary marker and found rates for *ascl1b, olig2* and *neurod1* of 100%, 10% and 20%, respectively (Table 1). F1 embryos positive for EGFP expression were tested for on-target precise integration at the genomic target site by PCR amplification and sequencing of the 5’ and 3’ genome/cassette junctions (Figure 1; Supplementary Figure 2). We found F0 transmission of precise integration alleles through the germline was relatively high, with frequencies at *ascl1b, olig2* and *neurod1* of 33% (1/3), 5% (1/20) and 20% (1/5), respectively (Table 1). These results at 3 independent loci demonstrate 2A-Cre in-frame targeted integration alleles can be efficiently recovered after screening a minimum of 20 F0 adults.

Single F1 adults harboring precise integration alleles were outcrossed to recover F2 adults, and single F2 adults were used to establish *Tg*(*ascl1b-2A-Cre)*^*is75*^, *Tg*(*olig2-2A-Cre)*^*is76*^, and *Tg*(*neurod1-2A-Cre)*^*is77*^ F3 families. Because the 2A-Cre cassette was integrated early in the proneural gene coding sequence, a loss of function mutation should result; therefore, the transgenic lines are maintained as heterozygotes. The F3 heterozygous *ascl1b-2A-Cre, olig2-2A-Cre*, and *neurod1-2A-Cre* larvae and adults did not exhibit anatomical or behavioral phenotypes.

### Endogenous proneural 2A-Cre transgenic lines express Cre in a pattern that recapitulates the targeted gene

To confirm the *ascl1b-2A-Cre, olig2-2A-Cre*, and *neurod1-2A-Cre* integration lines express Cre in a pattern that recapitulates the targeted gene we used whole mount *in situ* hybridization to examine *Cre* transcript localization. Adult F2 or F3 heterozygotes were outcrossed to wild type WIK and 3 dpf sibling larvae were hybridized with a probe complementary to *Cre* or to the target gene mRNA. Each of the lines showed that the pattern of Cre expression was identical to the endogenous gene mRNA (Figure 2).

**Figure 2.**
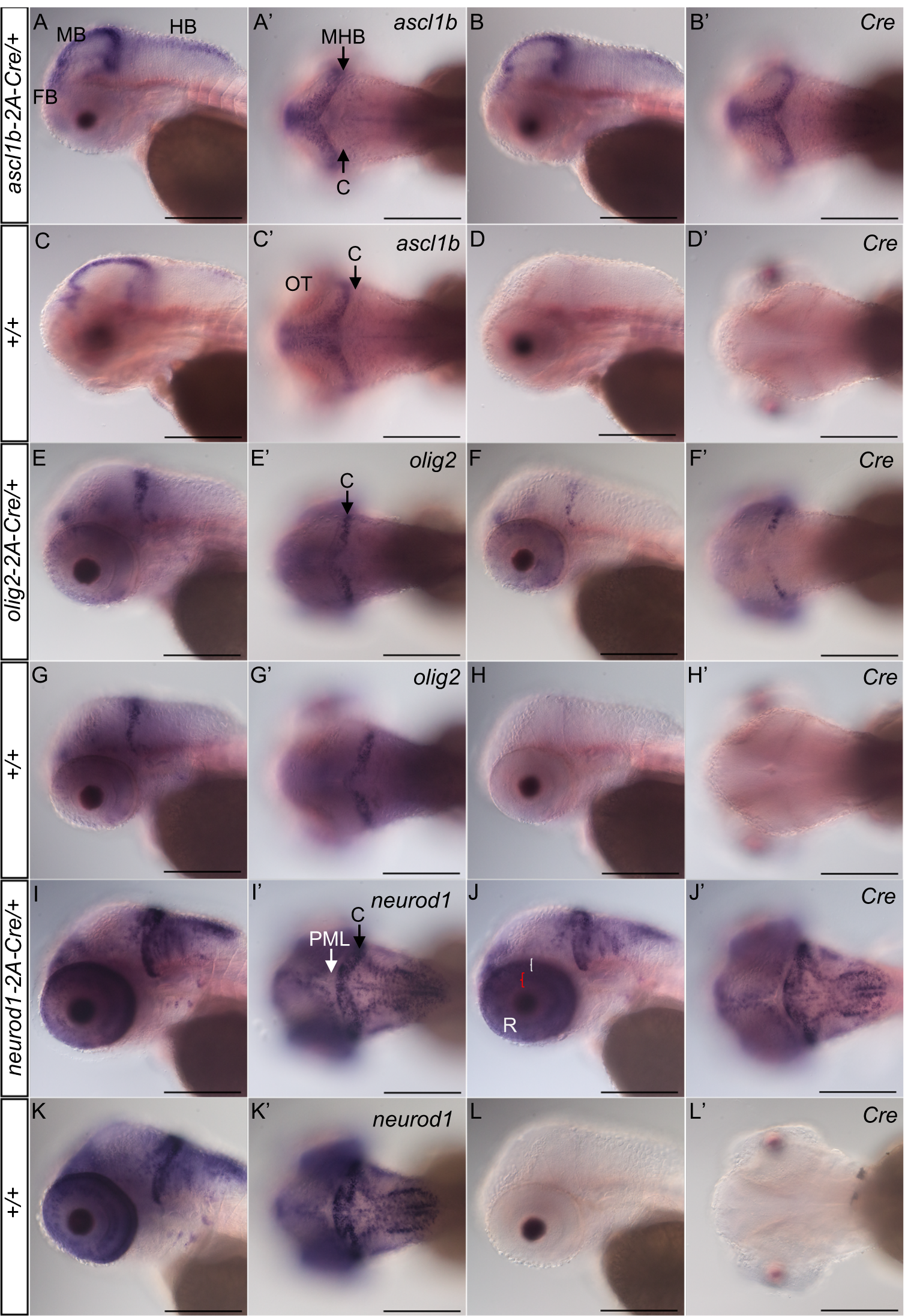
Expression of 2A-Cre integration alleles recapitulates *ascl1b, olig2*, and *neurod1* expression pattern. Whole mount *in situ* hybridization for endogenous genes and Cre was performed in 3 dpf larvae obtained from outcrossing *ascl1b-2A-Cre/+, olig2-2A-Cre/+*, and *neurod1-2A-Cre/+* lines to wild type WIK. Sibling larvae from the same clutch were sorted into lens EGFP-positive and -negative groups before fixation. For each genotype and probe 3 individual larvae were photographed. (A-D) *ascl1b* and Cre expression in *ascl1b-2A-Cre/+ larvae* show similar patterns were in the forebrain, and along the midbrain ventricle and midbrain-hindbrain border. (E-G) *olig2* and Cre expression in *olig2-2A-Cre/+* larvae were restricted to the forebrain, the posterior half of the cerebellum, and a subset of cells in the retina. (I-K) *neurod1* and Cre expression in *neurod1-2A-Cre/+* larvae were detected in the forebrain, adjacent to the midbrain and hindbrain ventricles and enriched in the cerebellum (I’, C black arrow), and in the retina inner (red bracket) and outer (white bracket) nuclear layers (J). *neurod1* and Cre expression were not detected in the posterior peripheral midbrain layer (I’, PML white arrow). Cre expression was not detected in wild type +/+ sibling larvae (D, H, L). *C*, cerebellum; *FB*, forebrain; *HB*, hindbrain; *MB*, midbrain; MHB, midbrain-hindbrain border; *NT*, neural tube; *PML*, peripheral midbrain layer; *R*, retina. Scale bar 250 μm.

*ascl1b-2A-Cre/+* larvae hybridized with *ascl1b* or *Cre* probes showed the same pattern of signal in the forebrain, along the midbrain ventricle and midbrain-hindbrain border in the optic tectum, and on the dorsal surface of the hindbrain (Figure 2 A-D), as previously reported (Wagle et al., 2004). *olig2-2A-Cre*/+ larvae showed *Cre* expression matched *olig2* in wild type larvae (Figure 2 E-H), (McFarland et al., 2008) and was restricted to the posterior half of the cerebellum, a subset of cells in the neural retina, and the ventral forebrain. Like the endogenous *neurod1* in wild type (Figure 2I, J) (Forbes-Osborne et al., 2013; Hozumi et al., 2012), *Cr*e expression in *neurod1-2A-Cre/+* was observed in the forebrain, adjacent to the midline in the midbrain and hindbrain, and absent from the peripheral midbrain layer of the optic tectum, consistent with *neuro*d1 expression in committed progenitors and newborn post-mitotic neurons. It was highly expressed in the cerebellum and in the inner and outer neural layers of the retina (Figure 2I-L). For all three lines, the consistency between the *Cre* and endogenous gene expression patterns suggested that both were controlled by the same regulatory elements, providing gene specific spatial and temporal Cre expression in neural progenitor populations defined by the proneural transcription factors.

### Endogenous neurogenic 2A-Cre transgenic lines lead to efficient Cre-*loxP* recombination in the expected neural cell populations defined by proneural gene expression

To confirm the stable *ascl1b-2A-Cre, olig2-2A-Cre*, and *neurod1-2A-Cre* driver lines express functional Cre recombinase in the expected neural progenitor cell populations, F0 adults were mated to the ubi:Switch line (Figure 3 A). The offspring were imaged at 3 dpf and showed expression had switched from EGFP to mCherry in neurons in the developing forebrain, midbrain, hindbrain and retinas of the double heterozygous 2A-Cre; ubi:switch larvae (Figure 3 B-D). *ascl1b-2A-Cre* led to near complete switching of EGFP to mCherry along the anterior-posterior axis of CNS, from the olfactory placode extending to the neural tube, with the exception of the neural retina (Figure 3 B). This is consistent with the early expression of *ascl1b* in the midbrain and hindbrain regions of the neural plate in 4-10 somite stage embryos, which broadens to cells in the anterior telencephalon, diencephalon, midbrain tegmentum and hindbrain by 25 somites/22 hours post fertilization, but is completely absent from the developing retina (Allende and Weinberg, 1994; Thisse and Thisse, 2005). Little switching occurred in cells outside of the CNS. These results indicated Cre expressed by from the *ascl1b-2A-Cre* allele led to efficient recombination at the *ubi:Switch* transgene *loxP* sites and excision of the floxed EGFP cassette.

**Figure 3.**
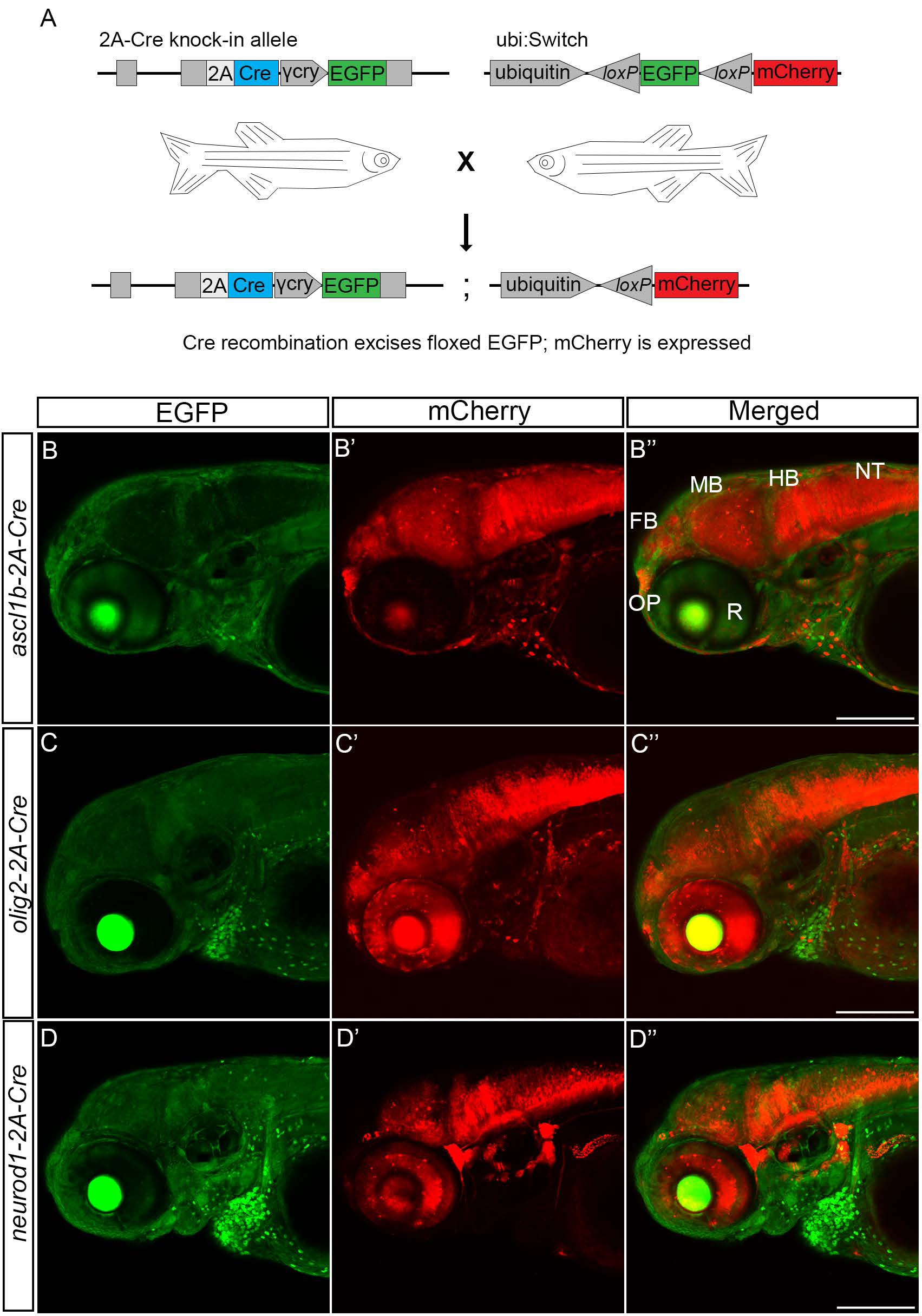
Transgenic *ascl1b-2A-Cre, olig2-2A-Cre* and *neurod1-2A-Cre* lines express functional Cre that promotes recombination at *loxP* sites in the expected neural progenitor populations. (A) F1 adults harboring precise integration alleles were mated to the recombination reporter line ubi:Switch to generate double transgenic 2A-Cre driver; ubi:Switch embryos. (B-D) Confocal imaging of 3 dpf double transgenic *ascl1b-2A-Cre*; *ubi:switch* (B), *olig2b-2A-Cre*; *ubi:switch* (C), and *neurod1-2A-Cre*; *ubi:switch* (D) larvae shows a switch from GFP to mCherry expression in neural cells derived from *ascl1b, olig2*, and *neurod1* progenitors. *FB*, forebrain; *γ-cry*, gamma crystallin promoter; *GFP*, green fluorescent protein; *HB*, hindbrain; *MB*, midbrain; *NT*, neural tube; *OP*, olfactory placode; *R*, retina. Scale bar 100 μm.

Similar to *ascl1b-2A-Cre*, the *olig2-2A-Cre* transgenic line led to recombination and switching from EGFP to mCherry expression throughout the CNS, including the retina (Figure 3 C). The pattern was consistent with early expression of *olig2* in the neural plate presumptive ventral diencephalon at 8 hpf (Park et al., 2002); forebrain dorsal thalamus, subpallium, posterior ventral thalamus and posterior tuberculum at 24-30 hpf (Borodovsky et al., 2009); and along the forebrain ventricles, retina, midbrain/hindbrain border and cerebellum at 48 hpf (Kani et al., 2010; Luo et al., 2016). The pattern of switching induced by the *neurod1-2A-Cre* line extended throughout neurons in the forebrain, midbrain, hindbrain, retina, and the PNS (Figure 3 D), as expected, given *neurod1* is expressed in committed neural progenitors and early post mitotic neurons throughout the nervous system (Rauch, 2003). Together with the *in situ* hybridization analyses, these results show the 2A-Cre knock-in lines express functional Cre recombinase in the pattern and cell type defined by the targeted proneural gene, leading to all progenitor cell descendants inheriting the recombined floxed allele.

In summary, in this study we applied our CRISPR/Cas9 short homology directed targeted integration strategy (Wierson et al., 2020) to generate a set of zebrafish proneural Cre drivers that express functional Cre recombinase in the pattern of the endogenous targeted *ascl1b, olig2*, and *neurod1* genes. Using 48 bp homology arms in the universal gRNA targeting vector we were able to efficiently recover precise in-frame 2A-Cre recombinase integration alleles after screening a maximum of only 20 adult F0 fish. The frequency of allele recovery were similar to our previous report describing efficient knock in of 2A-RFP and 2A-EGFP reporter cassettes at eight zebrafish loci (Wierson et al., 2020). Although *ascl1b, olig2*, and *neurod1* do not show haploinsufficiency, for other loci it may be beneficial to maintain the wild type endogenous gene expression in the targeted allele. Previous work showed homozygous *otx2:CreERT2* embryos generated by CreERT2 integration upstream of the zebrafish *otx2* initiating ATG were morphologically normal, but endogenous *otx2* expression was reduced (Kesavan et al., 2018). Nagy *et al*. (2019) inserted Cre before the translation termination codon in the mouse megakaryocyte/platelet-specific *Gp1ba* gene, which led to a decrease in GPIbα protein expression and multiple platelet defects. These studies suggest the potential for targeted integration to negatively impact endogenous gene expression will be specific to gene and location, and may require testing alternative design strategies. An alternative would be to target the termination codon with 2A-Cre as described by Nagy et al., (2019). Our efficient method for generating zebrafish transgenic Cre driver lines by CRISPR targeted integration can be readily applied to other loci, expanding opportunities to develop new recombinase genetic tools for investigation of specific cell types, developmental stages, or disease states.

## Experimental Procedures

### Zebrafish strains and maintenance

Zebrafish (*Danio rerio*) were maintained on an Aquatic Habitats (Pentair) or Aquaneering aquaculture system at 27^°^C on a 14 hour light/10 hour dark cycle. The wild type strain WIK was used to generate the knock in lines and was obtained from the Zebrafish International Resource Center (https://zebrafish.org/home/guide.php). The ubiquitously expressed floxed EGFP-mCherry expression line ubi:Switch (*Tg(ubi:loxP-EGFP-loxP-mCherry)*) was previously described (Mosimann *et al*., 2011). All zebrafish experiments were carried out in accordance with approved protocols from Iowa State University Animal Care and Use Committee Log#11-06-6252, in compliance with American Veterinary Medical Association and NIH guidelines for the humane use of animals in research.

### pPRISM-Cre vector and targeted integration

The pPRISM-Cre vector was designed to be compatible with our previously described short homology-based CRISPR/Cas9 knock in strategy (Wierson et al., 2020). The cassette in the pPRISM-Cre vector has two functional units: (1) a 2A-Cre cDNA cassette and (2) a secondary fluorescent reporter. The 2A-Cre unit is composed of a self-cleaving 2A peptide from porcine teschvirus-1, the Cre recombinase cDNA, and an ocean pout transcriptional termination and polyadenylation sequence (*Zoarces americanus*, Gibbs and Schmale, 2000). The viral 2A peptide has been used extensively in zebrafish for production of multicistronic messenger RNAs (Provost et al., 2007), allowing release of Cre protein from the short endogenous gene’s polypeptide. The secondary reporter cassette for visually track transgenic embryos contains the gamma crystallin (*γ-cry*) promoter, mini-intron, nls-EGFP, bovine growth hormone transcriptional termination and two SV40 polyadenylation sequences (Yang and Cvekl, 2005; Clark *et al*., 2011; J. H. Kim *et al*., 2011). Flanking the entire sequence are universal sgRNA (UgRNA) sites for Cas9 induced double strand breaks to liberate the short homology arms and drive integration of the Cre-reporter targeting cassette (Wierson et al., 2020).

CRISPR sgRNAs target sites in the coding sequence were selected and tested for efficient indel formation by co-injection of 25 pg sgRNA plus 300 pg Cas9 mRNA into 1 cell stage wild type WIK zebrafish embryos, followed by PCR amplification of the targeted exon and analysis of heteroduplex formation by gel electrophoresis. 48 bp 5’ and 3’ homology arms were designed and cloned into the pPRISM-Cre vector as described (Wierson et al., 2020). sgRNA and homology arm oligonucleotide sequences are listed in Supplementary Table 1. For targeted integration the injection mix contained 25 pg of genomic sgRNA, 25 pg of UgRNA, 10 pg of targeting vector, and 300 pg Cas9 mRNA. Wild type WIK zebrafish embryos at the 1-cell stage were injected with 2 nl of the injection mix and screened at 3 dpf for fluorescent secondary marker expression. All injected embryos showing EGFP expression in the lens were selected and raised to adulthood.

### Isolation of stable integration alleles and genome/vector junction analysis

To identify adult F0 founders, adult fish were outcrossed to wild type WIK and at least 75 embryos were screened for lens-specific EGFP expression. EGFP positive F1 embryos were selected for genome/vector junction analysis. Genomic DNA was extracted by digestion of single embryos in 50 mM NaOH at 95^°^C for 30 minutes and neutralized by addition of 1/10^th^ volume 1M Tris-HCl pH 8.0. Both 5’ and 3’ junctions were amplified by PCR using the primers listed in Supplementary Table 2, TOPO-TA cloned and sequenced. EGFP positive F1 embryos from transmitting founders were raised to adulthood and confirmed by fin clip. F1 adult animals with precise integration events were outcrossed to wild type WIK and F2 animals again screened for precise integration alleles. Single confirmed F2 adults were used to establish independent transgenic lines.

### *in situ* hybridization and live embryo imaging

Embryos were obtained from *ascl1b-2A-Cre/+, olig2-2A-Cre+* and *neurod1-2A-Cre/+* outcrossed to wild type WIK. Next, the collected embryos were placed in embryo media with 0.003% 1-phenyl 2-thiourea (PTU) at 24 hours post fertilization to block pigment production. At 3 dpf the PTU treated embryos were anesthetized in 160 ug/ml tricaine methane sulfonate and were fixed overnight at 4^°^C in 4% paraformaldehyde in PBS. Whole mount *in situ* hybridization was performed as described previously (McGrail et al., 2010). 3 dpf larvae were fixed overnight at 4^°^C in 4% paraformaldehyde or 4% paraformaldehyde/4% sucrose in PBS. *neurod1, olig2* and *ascl1b* cDNAs were cloned by RT-PCR using total RNA isolated from 3 dpf embryos and SuperScript III (Invitrogen). Primers for reverse transcription and PCR are listed in Supplementary Table 2. Digoxigenin-labeled probes were generated from linear plasmid DNA using DIG RNA Labeling Mix (Roche) and hybridized probes were detected with anti-digoxigenin antibody (Anti-Digoxigenin-AP, Roche) and NBT/BCIP (Roche). Larvae were imaged on a Zeiss Axioskop 2 and photographed with a Cannon Rebel T3 camera. Larvae for live imaging were placed in embryo media with 0.003% 1-phenyl 2-thiourea (PTU) at 24 hours post fertilization to block pigment production. At 2 or 3 dpf the PTU treated embryos were anesthetized in 160ug/ml tricaine methane sulfonate and mounted on slides in 1.2% low-melting agarose/160ug/ml tricaine methane sulfonate. Images were captured on a Zeiss LSM 700 laser scanning confocal microscope.

## Acknowledgements

The authors thank Dr. Leonard Zon (Harvard University) for the *Tg(ubi:loxP-EGFP-loxP-mCherry*) ubi:Switch transgenic zebrafish line. This study was supported by the Office of The Director of the National Institutes of Health under Award Number R24OD020166 (MM, JJE, KJC, SCE), and by a graduate scholarship from the CNPq Brazilian National Council for Scientific and Technological Development (MPA).

## Figure Legends

**Supplementary Figure 1.**
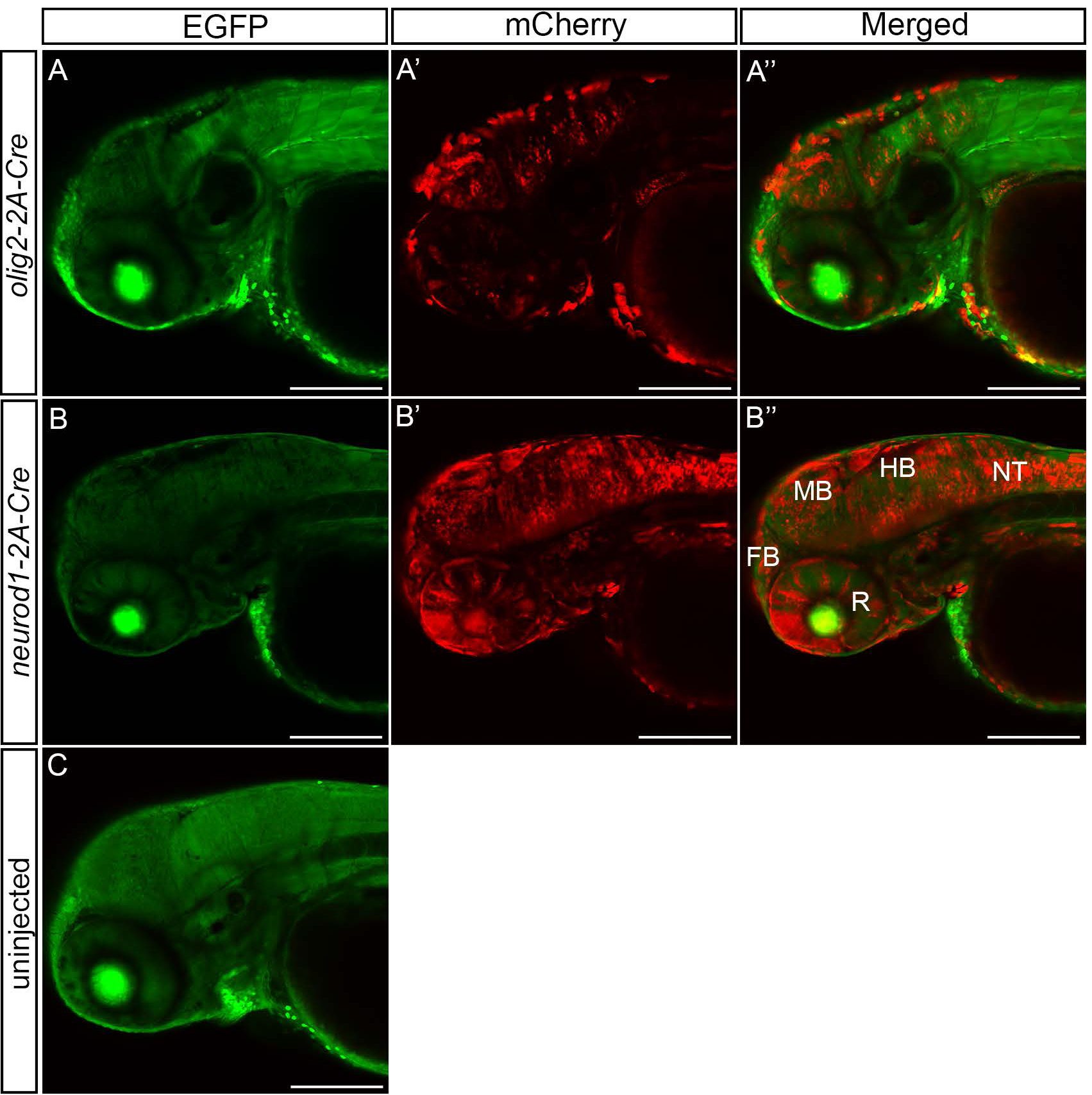
CRISPR/Cas9 targeted integration of *olig2-2A-Cre* and *neurod1-2A-Cre* cassettes leads to efficient Cre-*loxP* recombination in F0 mosaic animals. (A) 3 dpf *ubi:Switch* embryos that were injected with Cas9 mRNA, *olig2* exon 2 sgRNA, the universal short guide RNA UgRNA, and *olig2-2A-Cre* donor vector show a switch from GFP to mCherry expression in the midbrain, hindbrain, and neural tube, indicating Cre-mediated excision of the floxed GFP cassette. (B) Embryos injected with Cas9 mRNA, *neurod1* exon 2 sgRNA, the universal short guide RNA UgRNA, and the *neurod1-2A-Cre* targeting vector show the GFP to mCherry switch in presumed neurons throughout the entire developing central nervous system, including the retina. (C) Un-injected animals express only GFP. *GFP*, green fluorescent protein; *FB*, forebrain; *HB*, hindbrain; *MB*, midbrain; *NT*, neural tube; *R*, retina. Scale bar 100 μm.

**Supplementary Figure 2.**
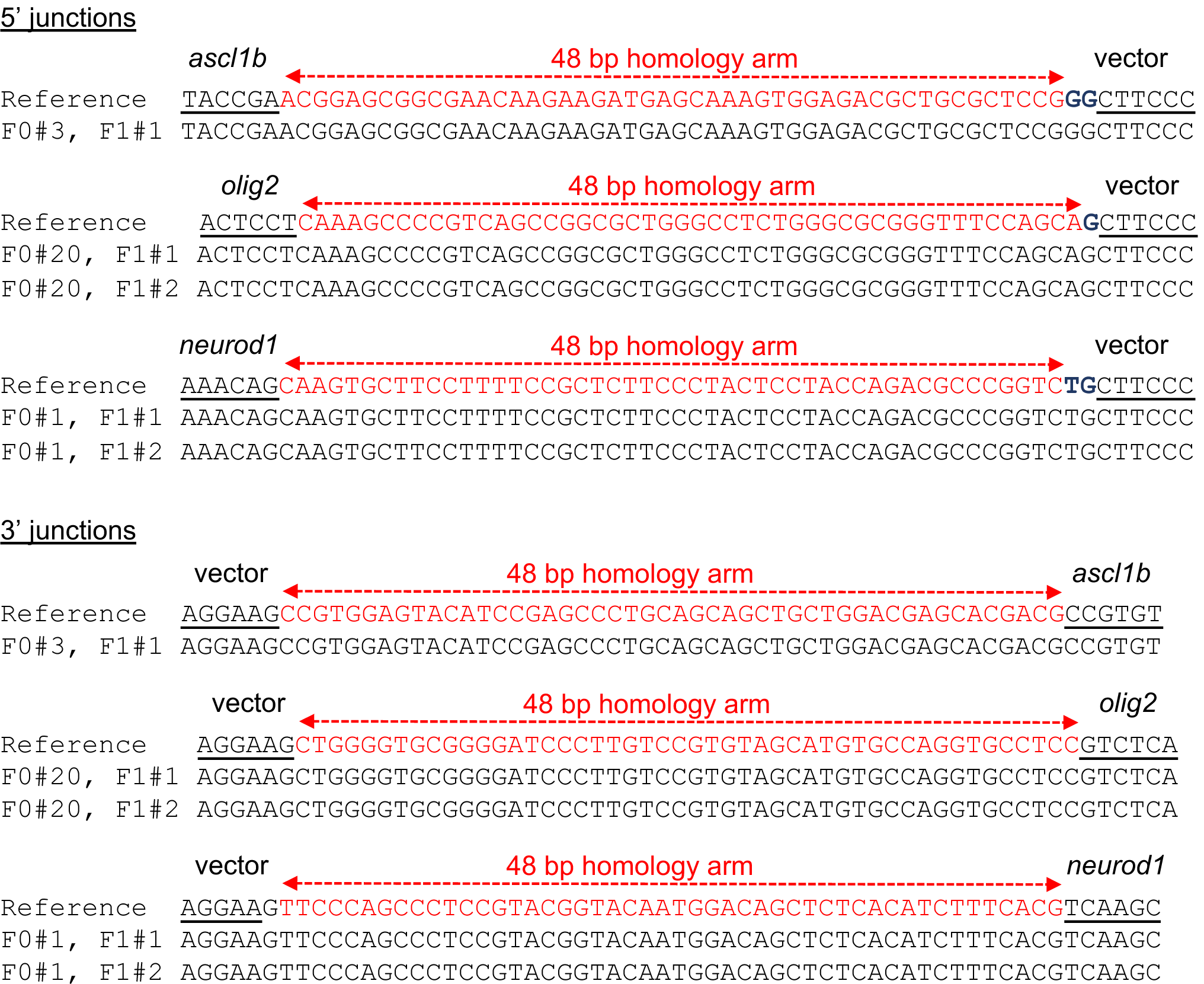
Sequence analysis of genome/vector junctions in F1 *ascl1b-2A-Cre, olig2-2A-Cre* and *neurod1-2a-Cre* transgenic F1 adults. 5’ and 3’ genome/vector junctions were PCR amplified, TA cloned and sequenced. Sequences were aligned to the reference sequence expected following a precise integration event. Red nucleotides represent the 48 bp homology arms used in the targeting vector. Blue letters represent additional nucleotides used to maintain the reading frame between the targeted exon and 2A-Cre cassette.

**Supplementary Table 1.**
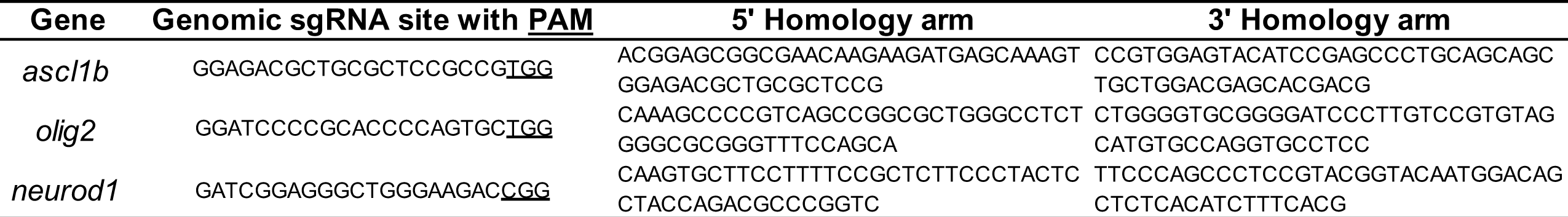
CRISPR sgRNA target sites and vector homology arm sequences

**Supplementary Table 2.**
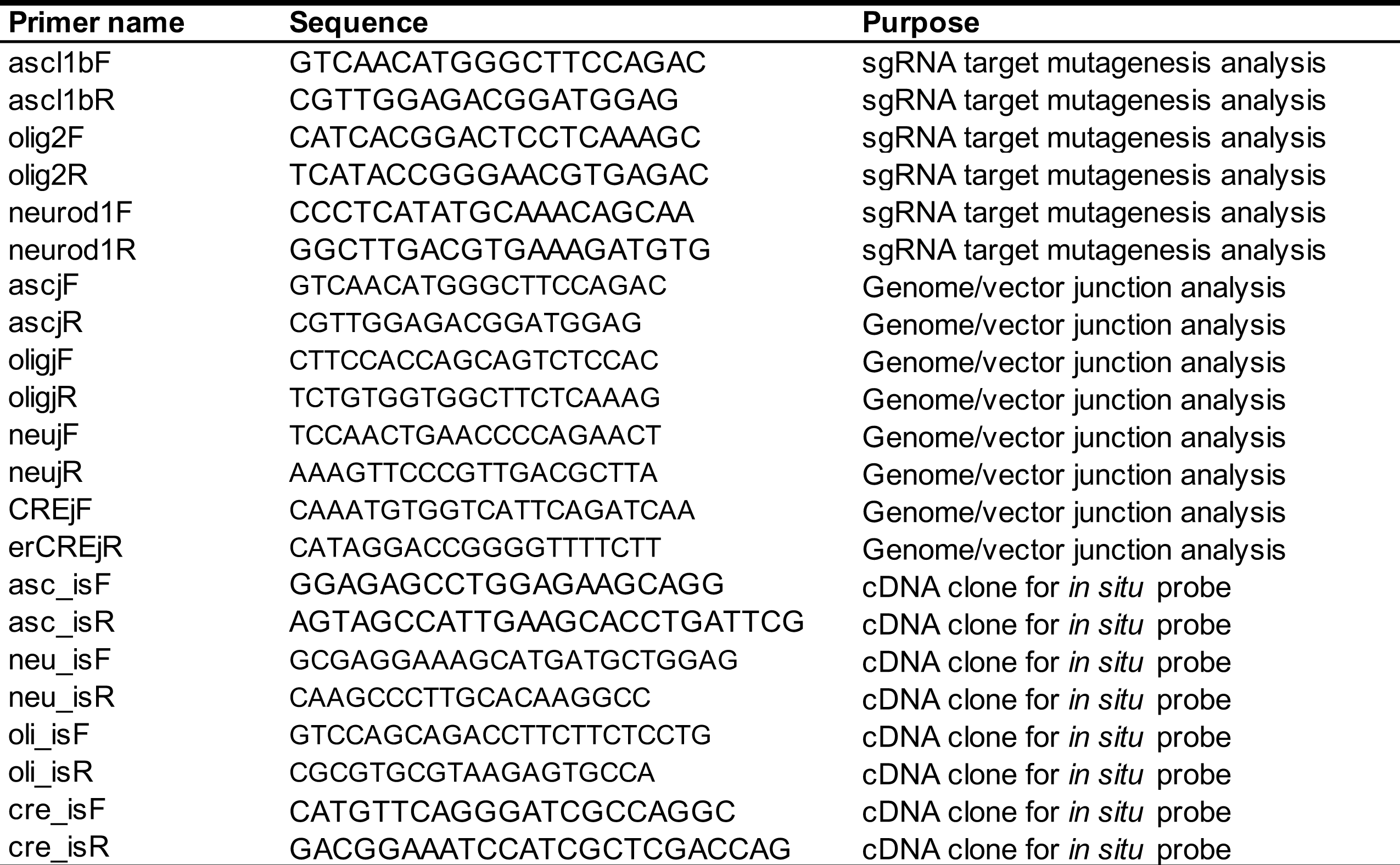
Primer sequences

